# Pharmacology of W-18 and W-15

**DOI:** 10.1101/065623

**Authors:** Xi-Ping Huang, Tao Che, Thomas J Mangano, Valerie Le Rouzic, Ying-Xian Pan, Susruta Majumdar, Michael Cameron, Michael Bauman, Gavril W. Pasternak, Bryan L Roth

**Author notes:** Correspondence to: Bryan L. Roth MD, PhD 4072 Genetic Medicine Building, Department of Pharmacology, UNC School of Medicine, Chapel Hill, NC 27514.

## Abstract

W-18 (1-(4-Nitrophenylethyl)piperidylidene-2-(4-chlorophenyl)sulfonamide)and W-15 (4-chloro-N-[1-(2-phenylethyl)-2-piperidinylidene]-benzenesulfonamide) represent two emerging drugs of abuse chemically related to the potent opioid agonist fentanyl (N-(1-(2-phenylethyl)-4-piperidinyl)-N-phenylpropanamide). Here we describe the comprehensive pharmacological profiles of W-18 and W-15. Although W-18 and W-15 have been described as having potent anti-nociceptive activity and are presumed to interact with opioid receptors, we found them to be without detectible opioid activity at μ, δ, κ and nociception opioid receptors in a variety of assays. We also tested W-18 and W-15 for activity as allosteric modulators at opioid receptors and found them devoid of significant positive or negative allosteric modulatory activity. Comprehensive profiling at essentially all the druggable G-protein coupled receptors in the human genome using the PRESTO-Tango platform revealed no significant activity. In silico predictions using the Similarity Ensemble Approach suggested activity for W-18 only weakly at H3-histamine receptors, which was not confirmed in radioligand binding studies. Weak activity at the sigma receptors and the peripheral benzodiazepine receptor were found for W-18 (K_i_=271 nM); W-15 displayed weak antagonist activity at 5-HT2-family serotonin receptors. W-18 is extensively metabolized, but its metabolites also lack opioid activity. W-18 and W-15 did inhibit hERG binding suggesting possible cardiovascular side-effects with high doses. Thus although W-18 and W-15 have been suggested to be potent opioid agonists, our results reveal no significant activity at these or other known targets for psychoactive drugs.

## INTRODUCTION

W-18 (1-(4-Nitrophenylethyl)piperidylidene-2-(4-chlorophenyl)sulfonamide) and W-15 (4-chloro-N-[1-(2-phenylethyl)-2-piperidinylidene]-benzenesulfonamide) were originally identified in the patent literature as analogues of the potent opioid agonist fentanyl (*N*-(1-(2-phenylethyl)-4-piperidinyl)-*N*-phenylpropanamide; (Fig 1) ^1^. In the original description, W-18 and W-15 (along with several other analogues) were described to be extraordinarily potent at inhibiting phenylquinone-induced writhing—a mouse model useful for assessing potential analgesic actions for a wide variety of drugs such as aspirin, antihistamines, opioids, antidepressants, sympathomimetics, actetylcholinesterase inhibitors and psychotomimetic opioids such as cyclazocine ^2^. Thus the inhibition of phenylquinone-induced writhing test is a relatively non-specific test for drugs with potential analgesic and other activities. In the original report there was no evidence that the activity in this model by either W-18 or W-15 was antagonized by the prototypical opioid antagonist naloxone although the related compound W-20 (1-cyclopropylmethylpiperidylidene-2-(4-chlorophenyl)sunfonamide) was reported to have modest naloxone-like activity.

**Figure 1.**
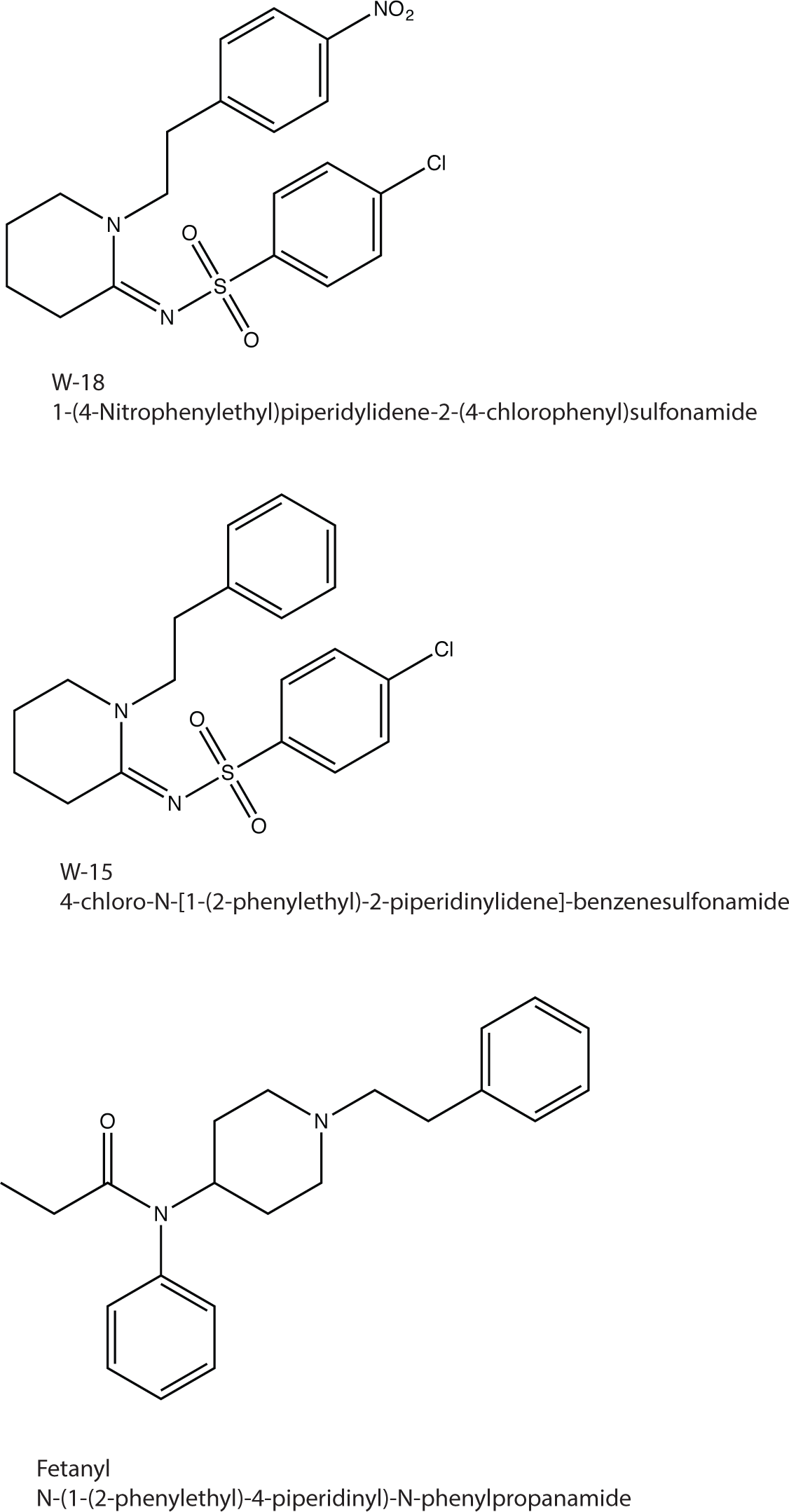
Structures of W-18, W-15 in comparison with fentanyl. As can be seen W-18 and W15 are 2-phenylethyl-2-piperidinyl compounds while fentanyl is a 2-pheylethyl-4-piperidinyl compound.

Recently W-18 ^3^, ^4^ ^5^ and perhaps W-15 have emerged as potential drug of abuse; W-18 is scheduled as a controlled substance in Canada^6^ and is being considered for inclusion in Schedule 1 in the U.S. To our knowledge there are no other published reports on the pharmacology of W-18 or W-15. Accordingly we here report a comprehensive evaluation of the in vitro molecular pharmacology of W-18 and W-15.

## METHODS

### Drugs

W-18 and W-15 were purchased from Cayman Chemicals (1180 East Ellsworth Road, Ann Arbor, MI 48108) and supplied as >98% pure. Independent LC-MS quality control revealed the compounds were both pure and authentic.

### Radioligand binding assays

Radioligand binding studies performed as previously described using cloned, human receptors ^7^ Comprehensive profiling was performed using the resources of the National Institute of Mental Health Psychoactive Drug Screening Program (NIMH-PDSP) as described ^7^. Initial screens were performed at 10,000 nM in quadruplicate. For targets with inhibition > 50%, we also carried out concentration-response binding assays to determine affinities (K_i_). Follow up assays on murine opioid receptors expressed in Chinese Hampster Ovary (CHO) cells were carried out as previously described^8^ ^9^.

### Metabolism studies

Metabolism was assessed using both mouse liver microsomes (MLM) and pooled human liver microsomes (HLM) incubated with NADPH (1 mM). Heat inactivated microsomes and incubations without NADPH were used as controls. At various time points 2-times volume of acetonitrile was added to stop the incubation and precipitate the protein so the sample could be analyzed by mass spectrometry. Samples were analyzed using multiple methods on a Sciex 6500 operating in multiple reaction monitoring mode using predicted mass transitions, using precursor ion and product ion scans. Samples were further analyzed on Thermo Scientific Q Exactive high resolution mass spectrometer using full scan mode operating at 70,000 resolution. Samples were also evaluated by HPLC using UV detection. HPLC-UV samples were processed by solid phase extraction, dried using a vacuum centrifuge and reconstituted in a minimal volume of mobile phase.

In addition, a W-18 mixed-metabolite sample was prepared by incubating 40 µl of a 20 mM W-18 DMSO stock with 5 ml of 1 mg/ml HLM or MLM and 1 mM NADPH for 90 minutes. Metabolites were isolated using solid phase extraction and dried using a speed vac. The mixed metabolite sample was re-solubilized and the pooled metabolites examined in opioid receptor binding assays using the murine receptors expressed in CHO cells at concentrations corresponding to an initial concentration of W-18 prior to the microsomes of 1 µM.

### Opioid receptor functional assays

Assays for agonist and antagonist activity at cloned opioid receptors were performed as previously described using cloned, human κ ^10^ ^11^. μ^12^, δ, and nociceptin opioid receptors. ^13^ ^14^. Assays for evaluating the allosteric modulation of opioid receptors were performed using graded concentrations of potential allosteric modulators in the absence and presence of increasing concentrations of the orthosteric ligand in a manner similar to that recently described by us ^15^.

### Profiling of W-18 and W-15 against the druggable GPCR-ome

These studies were performed using our recently described PRESTO-Tango resource ^16^, which allows for the unbiased interrogation of drugs against the entire, druggable GPCR-ome.

### Predictive pharmacology

We used the Similarity Ensemble Approach ^17^-^19^, essentially as previously described ^17, 18, 20^ to predict potential molecular targets for W-18 and W-15.

### Data integrity and reproducibility

All dose-response assays were performed several times and replicated independently while the GPCR-ome analysis was performed twice in quadruplicate at different concentrations.

### >RESULTS

#### Radioligand binding assays reveal no activity of W-18 at human, cloned opioid receptors

In our initial studies we evaluated the ability of W-15 and W-18 to interact with three canonical opioid receptors via radioligand binding studies performed as previously described using cloned, human ^10^ ^11^. ^12^, and δ opioid receptors (KOR, MOR and DOR, respectively). ^13^ ^14^. No significant inhibition of radioligand binding was measured up to concentrations as high as 10,000 nM (Table 1). These studies were followed by investigation of effects against the murine µ, κ and δ opioid receptors, with similar results. No inhibition of binding with concentrations as high as 1 µM were observed. W-15 and/or W-18 inhibited radioligand binding at several other receptors with low affinities, including the 5-HT_2A_, 5-HT_2B_, 5-HT_2C_ and 5-HT_6_ serotonin receptors benzodizepine receptors (BZP and PBR) (Table 1) and other miscellaneous targets with low affinities (Table 2). W-18 and W-15 had some activity at hERG suggestive of potential arrythmogenic activities at high doses (tables 1 and 2) ^21^.

**Table 1.**
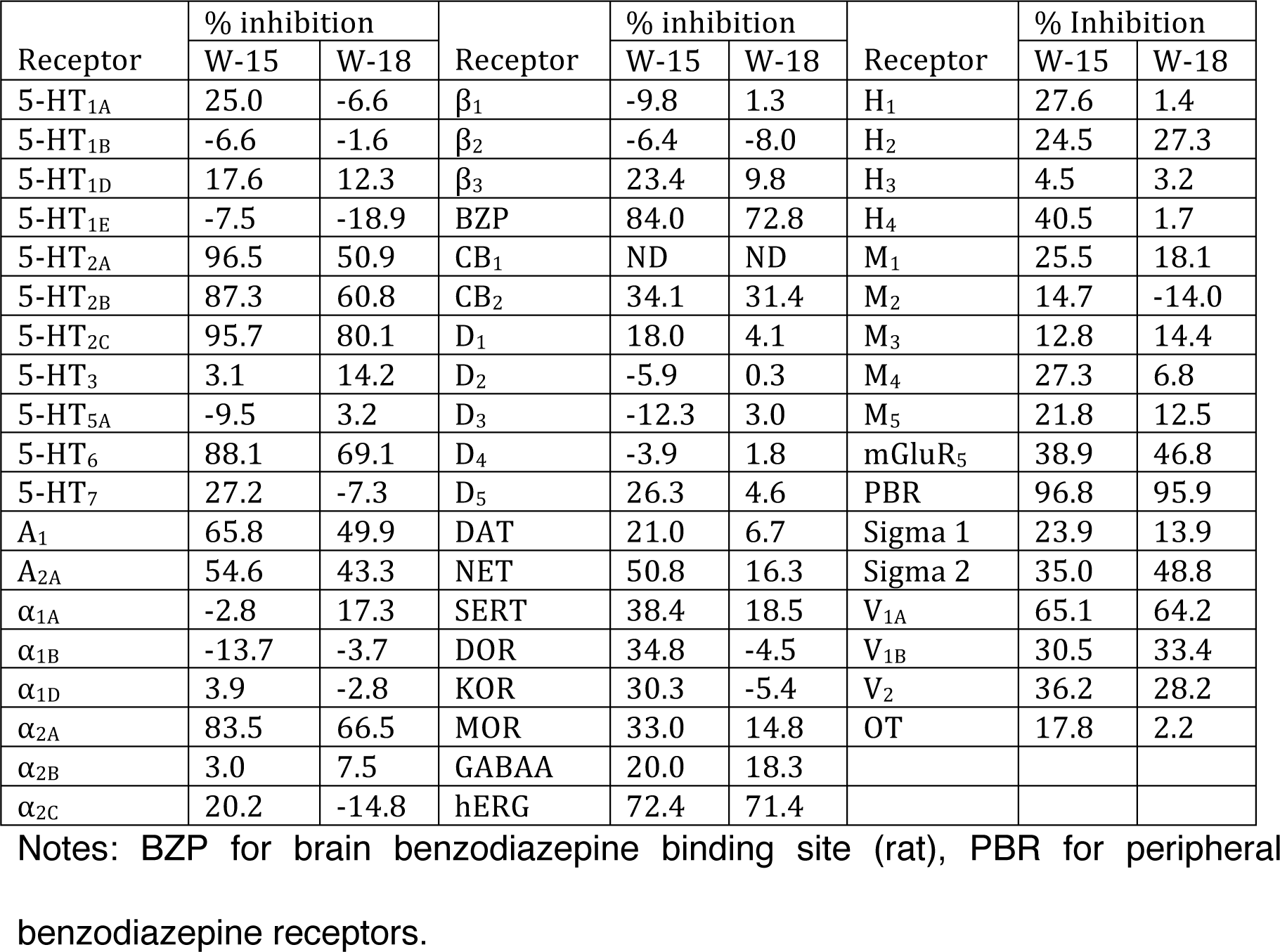
W-15 and W-18 binding profiles. Results represent percentage of inhibition at 10 µM. Values are averages from triplicate or quadruplicate set.

**Table 2.** Binding affinities for W-15 and W-18. Data represent mean of 2-4 separate determinations. Receptor

| Receptor | K<sub>i</sub> (pK<sub>i</sub> ± SEM, N) |  |
| --- | --- | --- |
|  | W-15 | W-18 |
| 5-HT<sub>2A</sub> | 109 nM (6.96 ± 0.05, 3) | 1751 nM ( 5.76 ± 0.20, 3) |
| 5-HT<sub>2B</sub> | 380 nM (6.42 ± 0.11, 3) | 2171 nM (5.66 ± 0.19, 3) |
| 5-HT<sub>2C</sub> | 52 nM (6.91 ± 0.07, 3) | 878 nM (6.06 ± 0.05, 3) |
| 5-HT<sub>6</sub> | 123 nM (6.91 ± 0.09, 3) | 1008 nM (6.00 ± 0.05, 3) |
| BZP | 933 nM (6.03 ± 0.06, 3 ) | 1023 nM (5.99 ± 0.55, 3 ) |
| PBR | 70 nM (7.16 ± 0.27, 4) | 77 nM (7.11 ± 0.24, 4) |
| A<sub>1</sub> | 3846 nM (5.42 ± 0.05, 2) | 5129 nM (5.29 ± 0.07, 2) |
| α<sub>2A</sub> | 2818 nM (5.55 ± 0.23, 4) | 2138 nM (5.67 ± 0.20, 4) |
| CB<sub>2</sub> | 8511 nM (5.07 ± 0.06, 3) | 7470 nM (5.13 ± 0.08, 3) |
| V<sub>1a</sub> | 3575 nM (5.45 ± 0.12, 3) | 3311 nM (5.48 ± 0.19, 3) |
| hERG | 2042 nM (5.69 ± 0.09, 3) | 1806 nM (5.74 ± 0.07, 3) |

#### Functional assays reveal no agonist, antagonist or allosteric modulator activity of W-18 or W-15 at human cloned opioid receptors

We next evaluated W-18 and W-15 for agonist and antagonist activity at cloned human opioid receptors in both G_i_-dependent inhibition of cAMP production and G-protein independent arrestin translocation assays (Figures 2 and 3). No significant agonist or antagonist activity was measured up to doses as high as 10,000 nM, although non-specific agonist effects were evident at high concentrations in these assays (Figs 2a-2d) We also evaluated the potential of W-18 to be allosteric modulators of opioid receptors and found no activities for either Gi or arrestin signaling (Figs 4-5).

**Figure 2.**
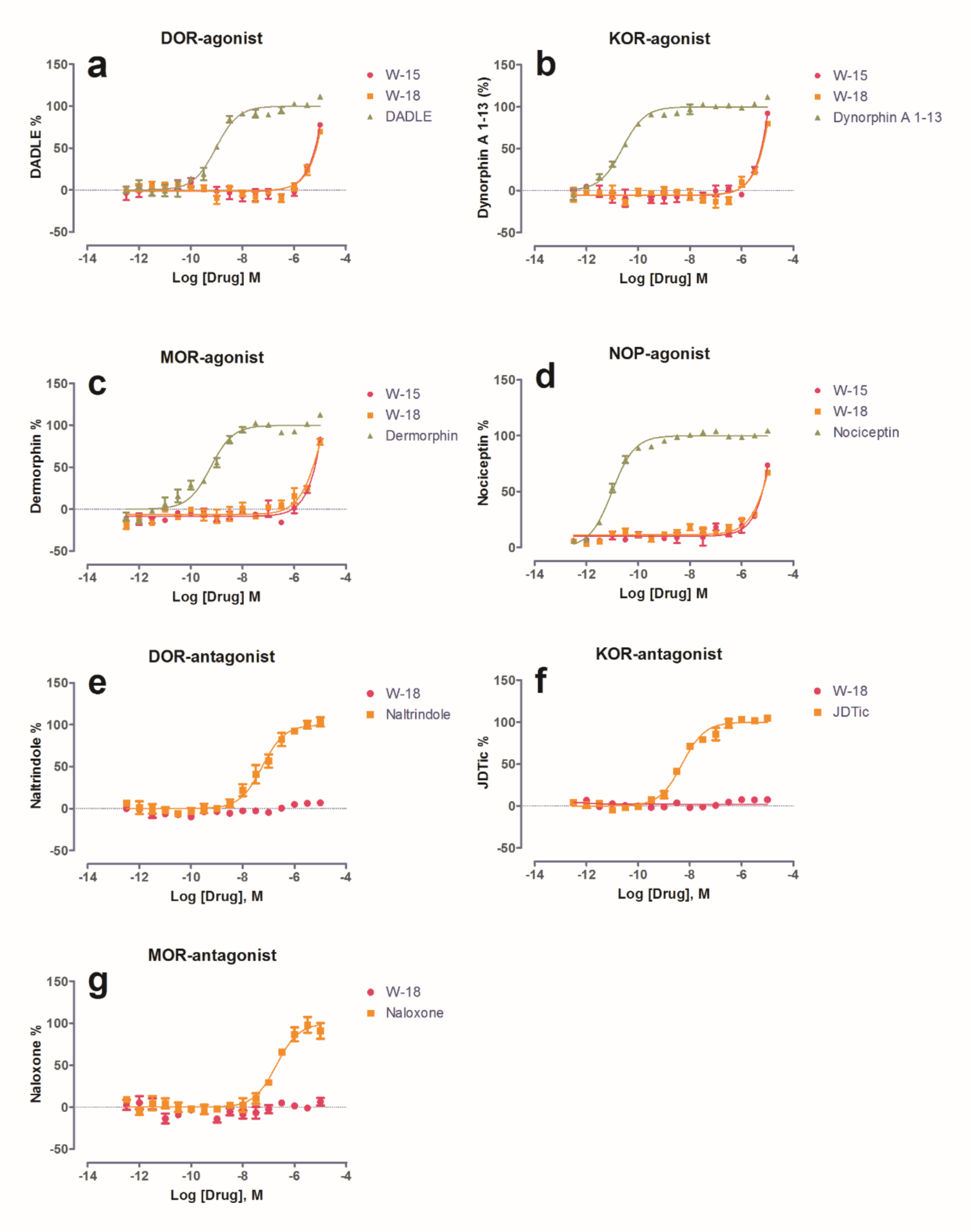
W-18 and W-15 lack activity at opioid receptors. Shown are data in which the effect of W-18 on (a) DOR, (b) KOR, (c) MOR and (d) NOP in agonist modes and (e) DOR, (f) KOR and (g) MOR in antagonist modes. As shown W-18 and W-15 have no substantial agonist or antagonist activities.

**FIGURE 3.**
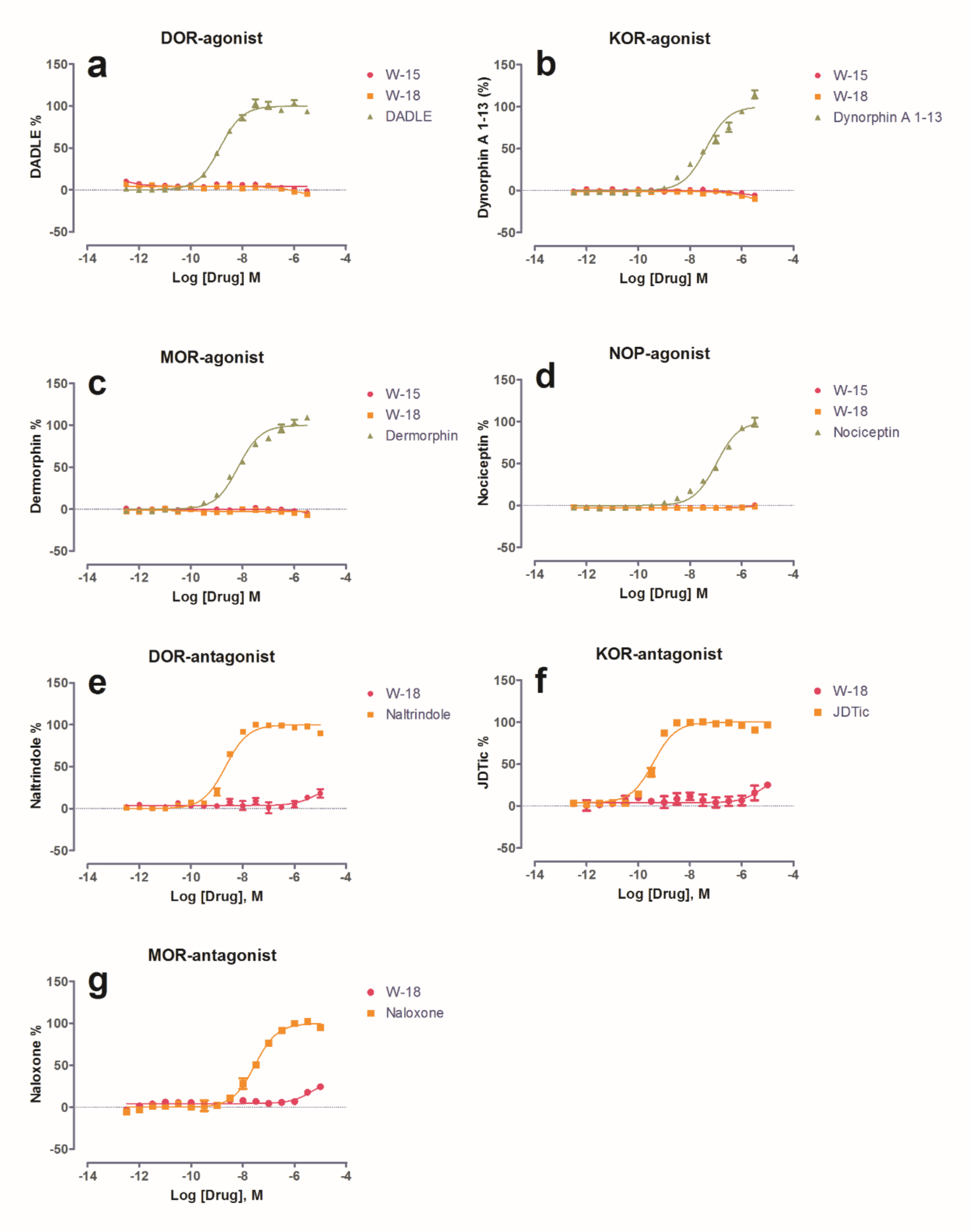
W-18 LACKS ACTIVITY FOR RECRUITING β-ARRESTIN2 IN BRET ASSAYS AT OPIOID RECEPTORS. Shown are representative graphs for MOR, KOR and DOR measuring agonist induced arrestin activity via bioluminescence energy transfer (BRET) quantified as previously detailed ^10^ in agonist (a-d) and antagonist (e-g) modes.

**FIGURE 4.**
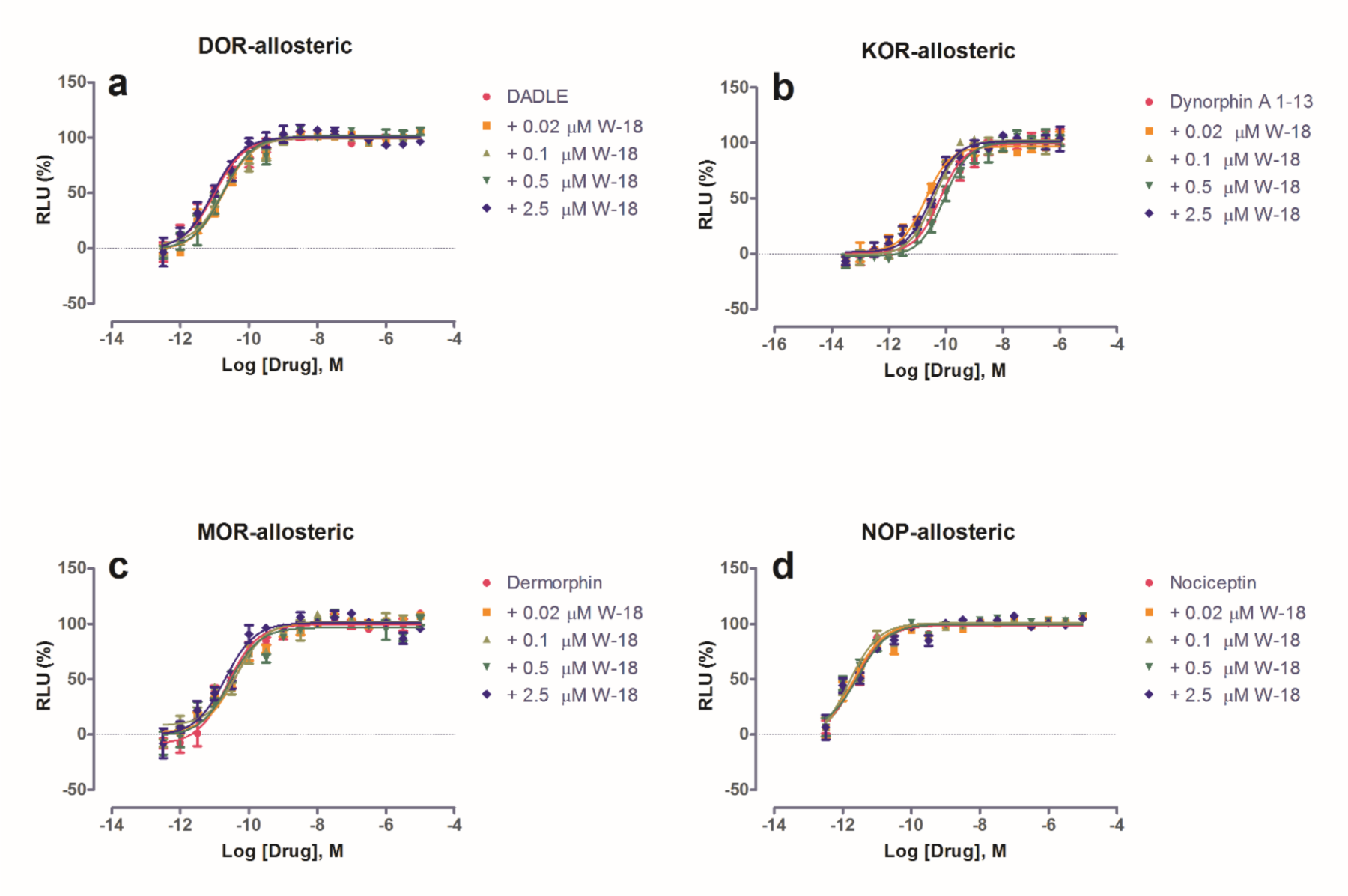
W-18 and W-15 lack activity as allosteric modulators of opioid receptors at G-protein signaling. Shown are data demonstrating lack of allosteric effect of W-18 and W-15 at opioid receptors for Gi signaling.

**Figure 5.**
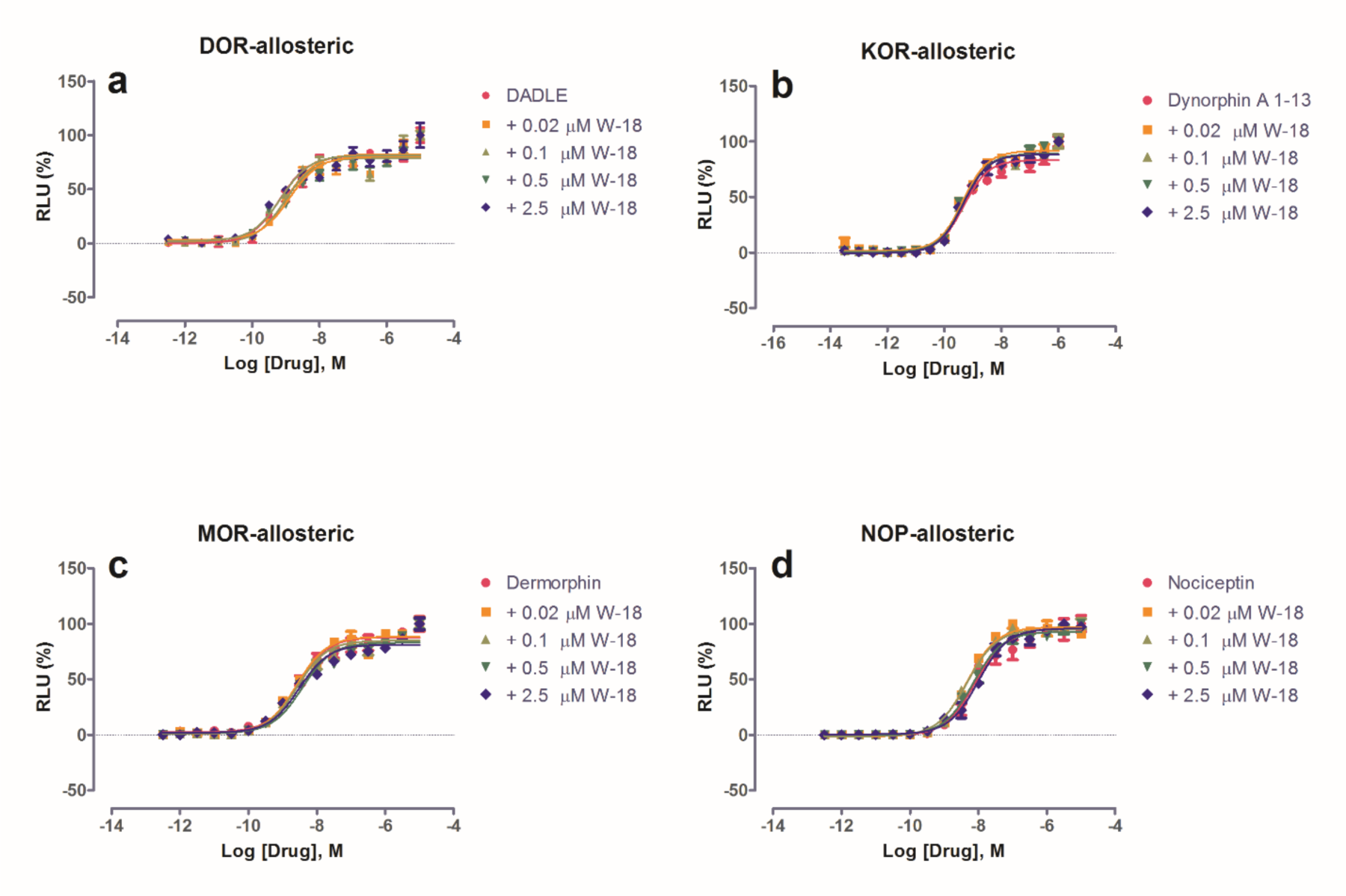
Lack of effect of W-18 and W-15 for allosterically modulating opioid activity at arrestin signaling. Shown are data demonstrating lack of allosteric effect of W-18 and W-15 at opioid receptors for arrestin translocation.

### Functional assays reveal antagonist activity at human 5-HT receptors

Since W-15 and W-18 displayed binding activities at 5-HT_2_ and 5-HT_6_ receptors, we examined their functional activity at 5-HT_2A_, 5-HT_2B_, and 5-HT_2C_ receptors for calcium mobilization and at 5-HT_6_ receptors for arrestin signaling. W-15 and W-18 showed no agonist activity and modest antagonist activity at 5-HT_2_ receptors (Fig 6). Schild analysis with W-15 indicated competitive antagonism against 5-HT at 5-HT_2_ receptors (Fig 7). At 5-HT_6_ receptors, W-15 and W-18 displayed no agonist activity (Fig 8). W-15 and W-18 also displayed weak binding at A_1_-adenosine, α_2A_-adrenergic, and CB_2_ cannabinoid receptors, we therefore examined the nature of their activity at these receptors. Our results (Fig 8) indicated that they had no agonist activity at any of these examined receptors.

**FIGURE 6.**
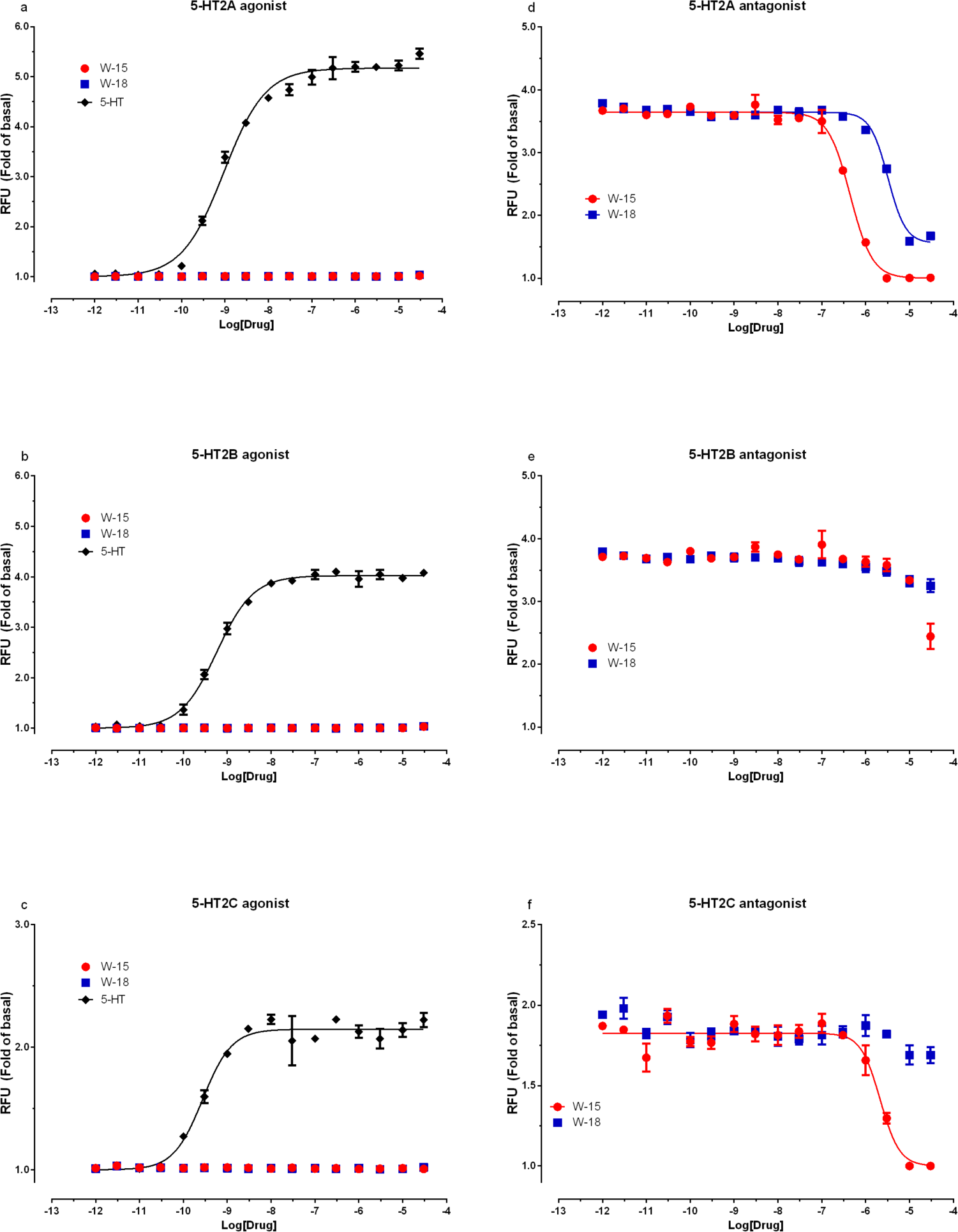
W-18 and W-15 display weak antagonist activities at 5-HT2-family receptors.

**FIGURE 7.**
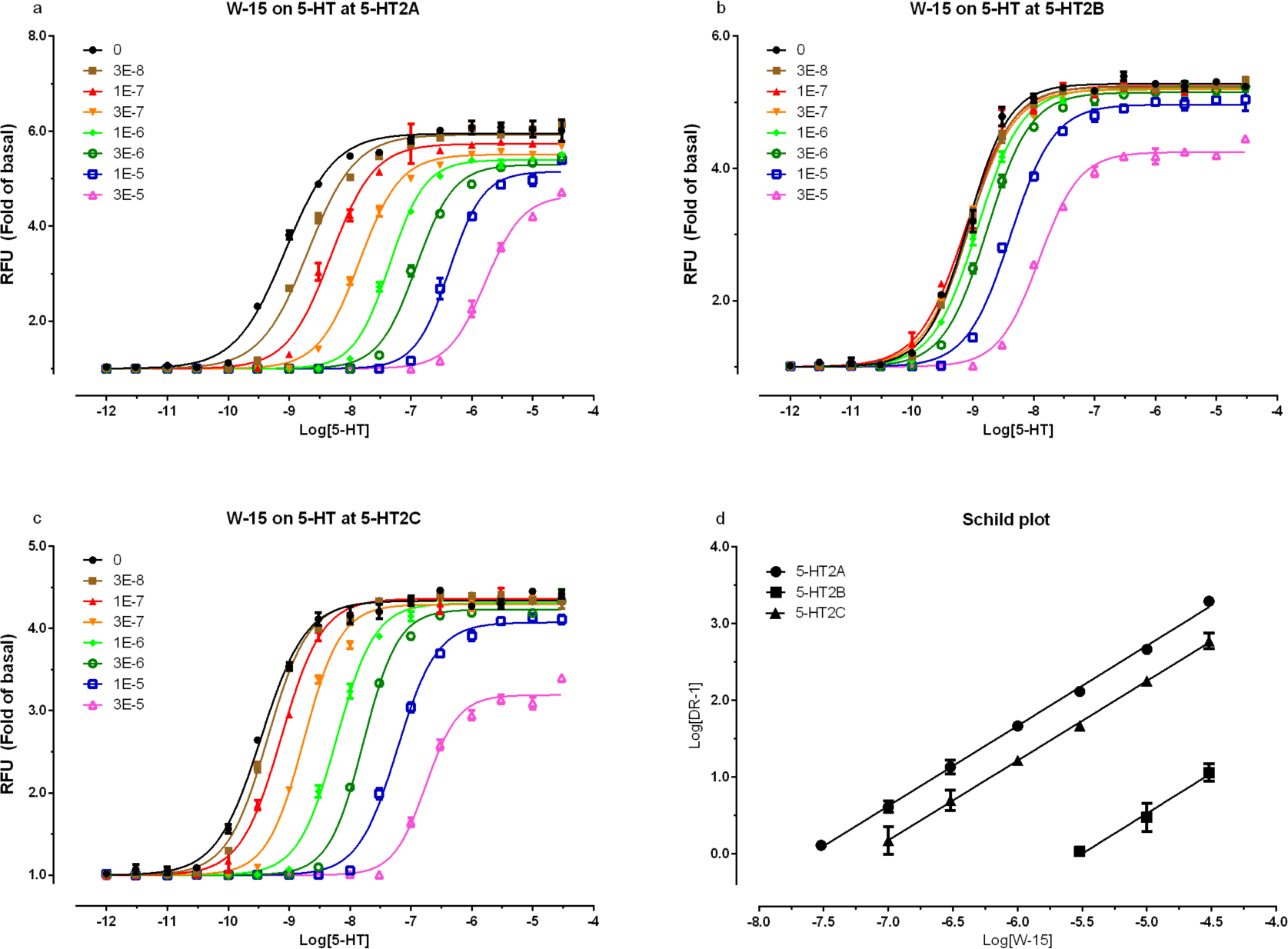
Schild analysis reveals W-15 is an orthosteric 5-HT_2A_, 5-HT_2B_ and 5-HT_2C_ antagonist.

**Figure 8.**
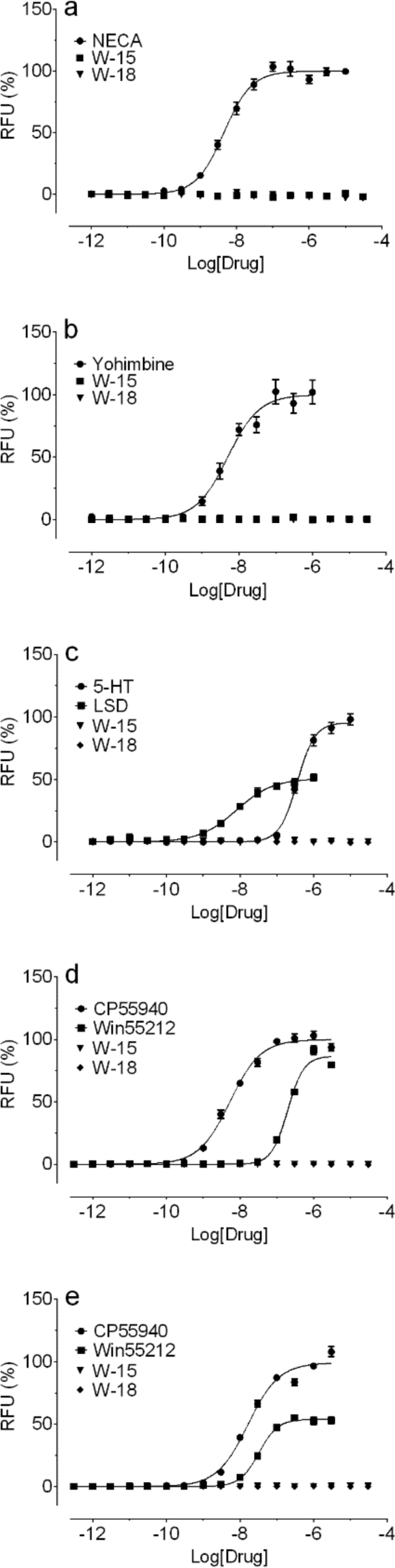
Lack of agonist activities of W-18 and W-15 at A_1_ (a), α_2A_ (b), 5-HT_6_ (c), CB_1_ (d), and CB_2_ (e), receptors. As can be seen W-18 and W-15 lack agonist activity at the tested receptors

**FIGURE 9.**
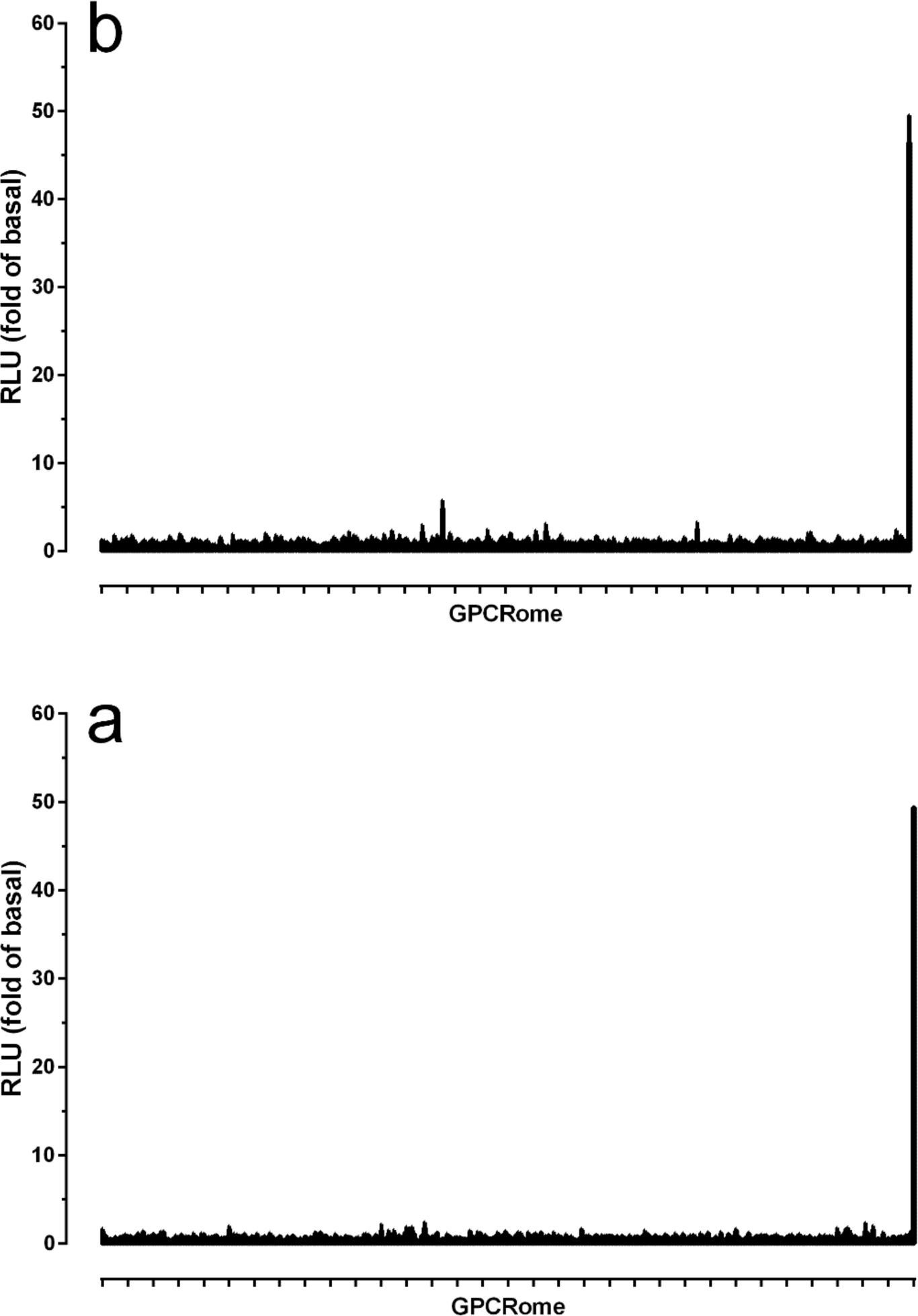
GPCR-OME PROFILING REVEALS NO SIGNIFICANT AGONIST ACTIVITY OF W-18 OR W-15. Shown are results in which the PRESTO-Tango resource was used to profile (a) W-18 and (b) W-15 against the 326 druggable GPCRs at 1 μM final concentration. No significant activity was found; the positive control represents quinpirole at the D2 dopamine receptor.

### Screening of W-18 and W-15 against the druggable GPCR-ome reveals no significant agonist activity at any human druggable GPCR

We next screened W-18 and W-15 against the druggable human GPCR-ome using our recently developed PRESTO-Tango resource ^16^. Although no significant agonist activity was reproducibly found, we did note that concentrations of W-18 higher than 1 μM tended to cause a decrease in the overall baseline activity for nearly every target consistent with a toxic activity at HEK cells with prolongued incubation (not shown).

We examined the metabolism of W-18. W-18 was extensively metabolized by both human and mouse liver microsomes (Supplemental Figs. 1-4), resulting in multiple mono and di-hydroxylated metabolites as well as a dealkylated and an amino metabolite from reduction of the nigro group. To determine if W-18 was a prodrug, releasing an active compound through metabolism, we incubated W-18 with liver microsomes, extracted the remaining W-18 and its metabolites and tested the mixture in opioid receptor binding assays at a concentration corresponding to 1 µM W-18 prior to the microsomal incubation. The mixture of W-18 and metabolites failed to significantly inhibit binding to murine µ, δ or κ opioid receptors expressed in CHO cells (not shown).

### >In silico studies

We also used the Similarity Ensemble Approach to determine if any molecular targets might be predicted with high expectancies to have activity for W-18. No target(s) emerged with high confidence although weak predictions (E-value=4.67 e^-1^) suggested activity at were for H3-histamine for W-18 which was unconfirmed (Table 1).

### In vivo evaluation

In the initial patent report W-18 was active in the writing assay, with a potenty many times that of morphine. when administred in mice at doses as high as 1 mg/kg, s.c., W-18 failed to demonstrate any activity in either the radiant heat tail flick assay or the writhing assay.no straub tail was seen. A general behavioral action of burrowing into the bedding was ovserved and the animals were clearly atypical in their overall behavior, showing a general tunneling behavior. Administration of naloxone failed to counteract these behaviors. similar burroeing behaviors were seen in rats after s.c administration of0.3 and 1,0mg/kg.

## DISCUSSION

The major findings of these studies are that both W-18 and W-15 are devoid of significant opioid receptor activity. Although W-18 and W-15 were described in the initial patent report as having potent analgesic activity in the phenylquinone-induced writhing assay, it should be noted that the test used cannot definitively identify the molecular target for the presumed ‘analgesic activity’. Thus as originally reported ^2^ a large assortment of pharmacological agents show activity in this assay including cholinesterase inhibitors, antihistamines and antidepressants ^22^ In our hands, W-18 revealed no antinoceptive activity in either the thermal radiant heat tail flick assay or the writhing assay. Furthermore, the atypical behaviors observed with the drugs (i.e. tunneling) failed to be reversed by naloxone, indicating that opioid receptors are not involved.

It is conceivable that W-18 and W-15 are prodrugs which require metabolic transformation for their effects in humans as has been seen for toxic pharmaceuticals including fenfluramine ^23^ and methysergide. ^24^ Even the prescribed analgesic codeine requires demethylation to generate morphine for activity in both receptor bniding assays and in vivo. W-18 undergoes extensive metabolism in both human and mouse liver microsomes, However, these metabolites do not reveal any affinity for opioid receptors either. Behaviorally, W-18 failed to elicit any naloxone-sensitive actions. Also, we saw no inhibition of murine opioid receptor binding from the W—18 metabolite mixture. Although further studies aimed at identifying the individual metabolite(s) and determining their pharmacological activities are needed and may uncover mechanisms contributing to the abuse potential of W-18, our studies strongly indicate that opioid systems are not involved.

In conclusion, comprehensive evaluation of the potential *in vitro* molecular pharmacology of W-18 and W-15 revealed no consensus targets for interaction. Although apparently devoid of opioid receptor activity, weak activity of W-18 at the peripheral benzodiazepine receptor was found and the drugs appear to exhibit toxicity in HEK cells with overnight incubation at concentrations higher than 1 μM. The literature has suggested that W-18 falls into the opioid class of drugs, possibly providing a false sense of security for those abusing the compound who may rely upon its reversibility by naloxone. Given the lack of apparent activity at human or mouse cloned opioid receptors for the parent compounds or the metabolites, the utility of naloxone and other opioid antagonists in treating over-dose in humans is unlikely to be successful. W-18 and W-15 had quite modest activity at hERG (Ki values >1 uM) suggesting that at high doses some potential for cardiac arrythmias is possible. This needs to be understood by both first responders and those abusing the compound.

## Acknowledgements

### ACKNOWLEDGEMENTS

This work was supported in part by the NIMH Psychoactive Drug Screening Program Contract and PO1DA035764 to BLR, grants from NIDA (DA06241) and from the Peter F. McManus Charitable Trust to GWP and a core grant from the NCI (CA08748) to MSKCC and an equipment grant 1S10OD010603 to MDC.

---

**SUPPLEMENTARY FIGURE 1:**
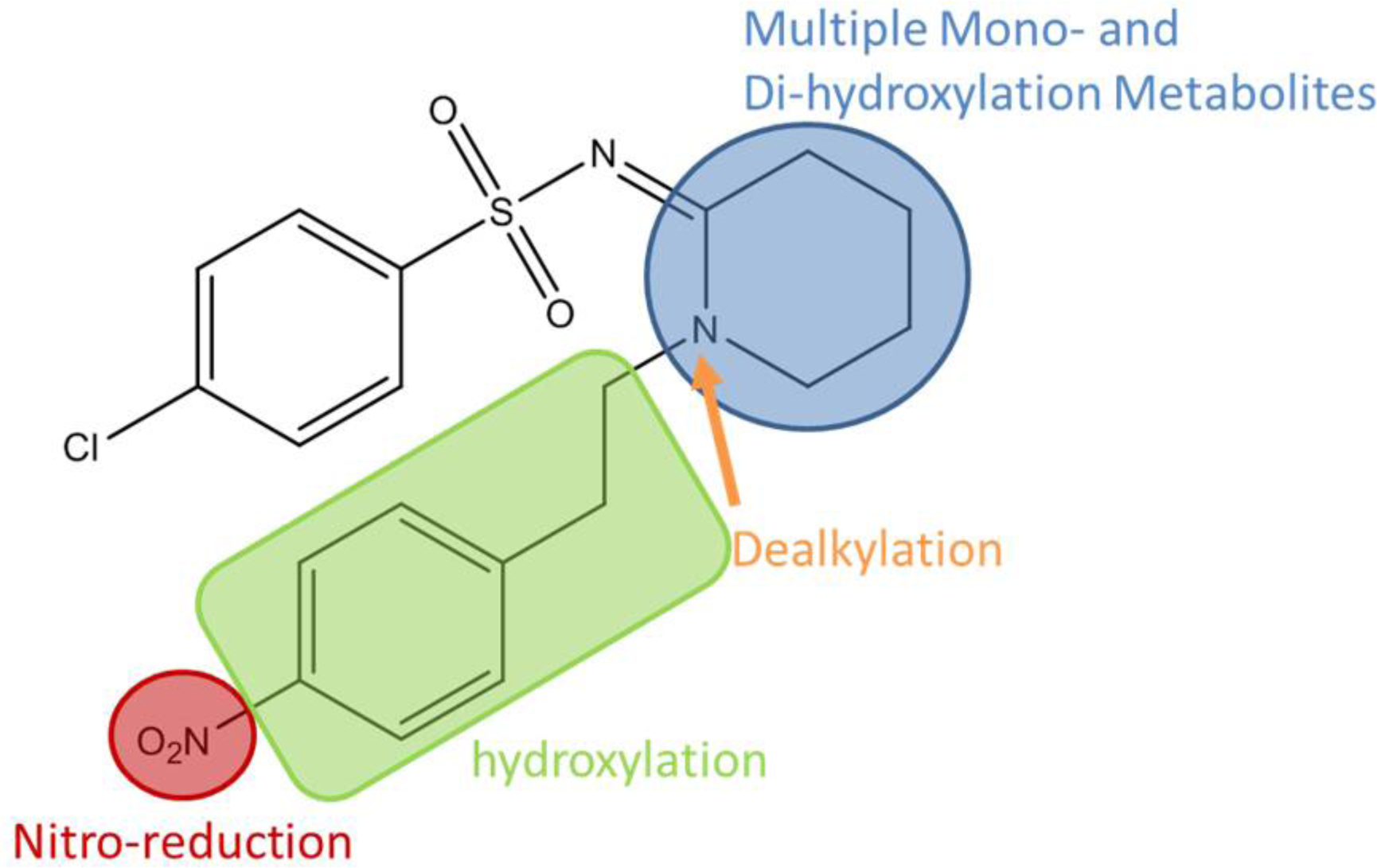
POTENTIAL SITES OF W-18 METABOLISM.

**Supplementary Fig 2A.**
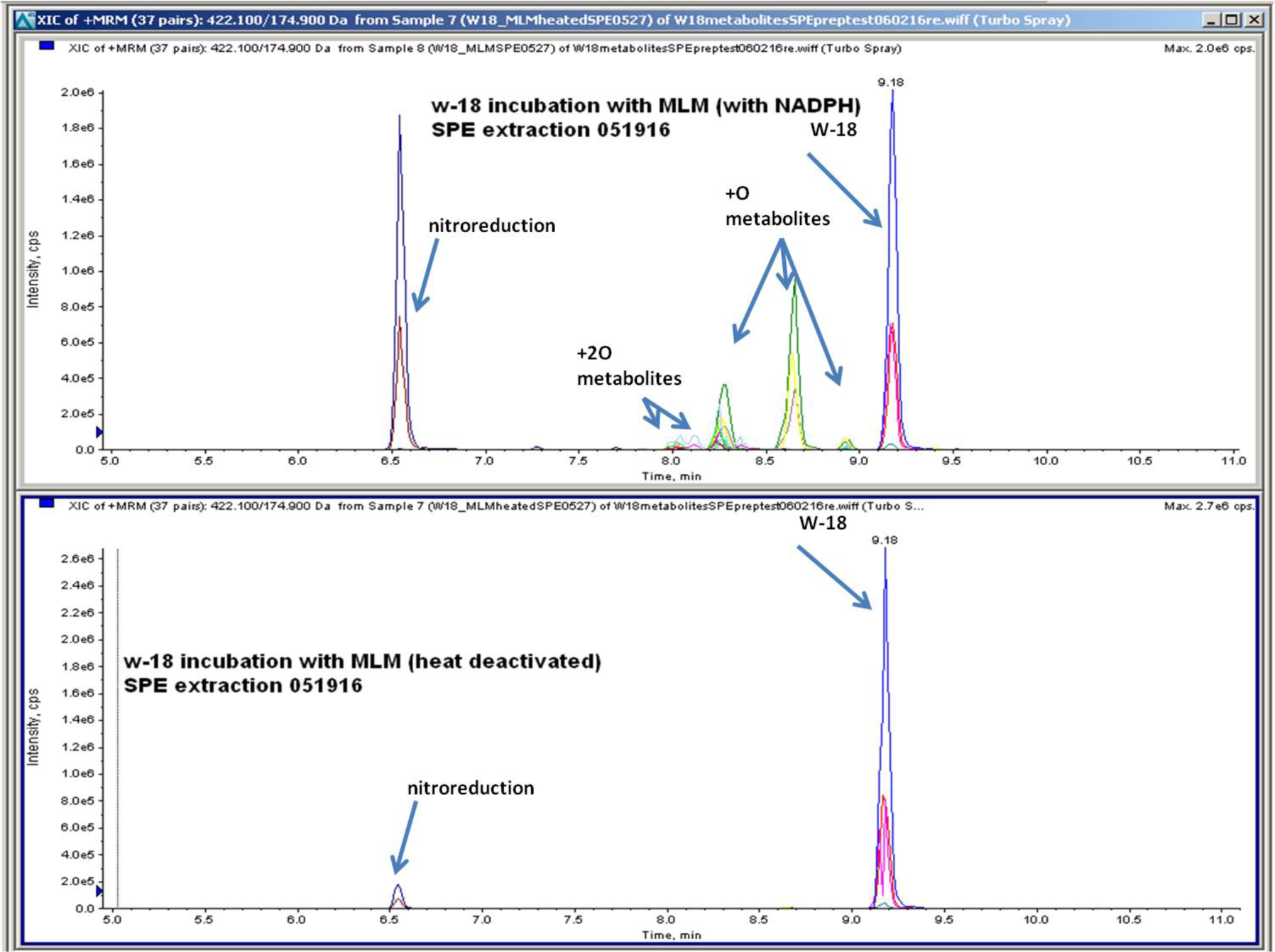
Metabolite profile of W-18 incubated with MLM and heat inactivated MLM (lower panel); MLM=mouse liver microsomes.

**Supplementary Figure 2A.**
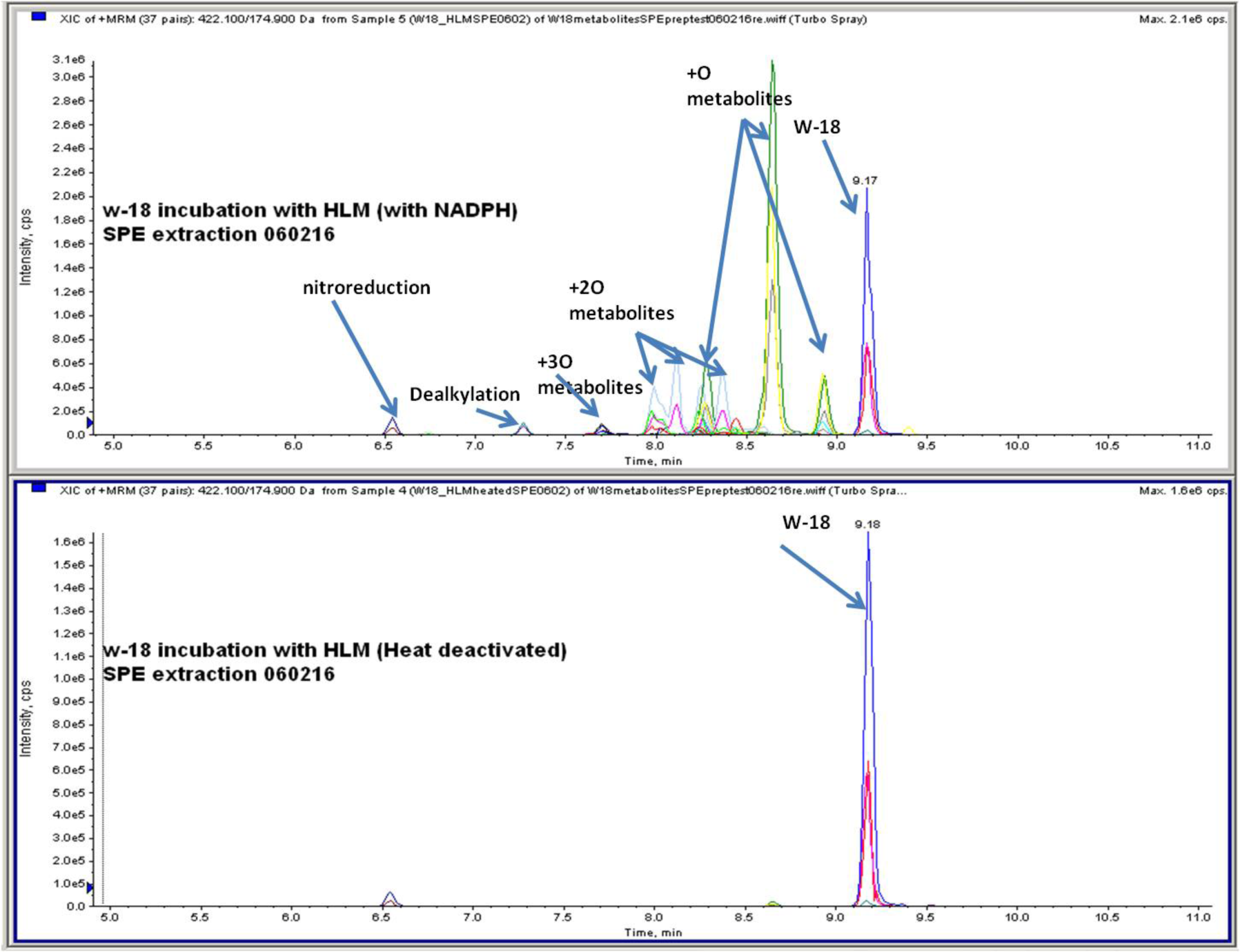
W-18 metabolite profile after incubation with HLM (upper) and heat-inactivated HLM. HLM=human liver microsomes.

**Supplementary Figure 3.**
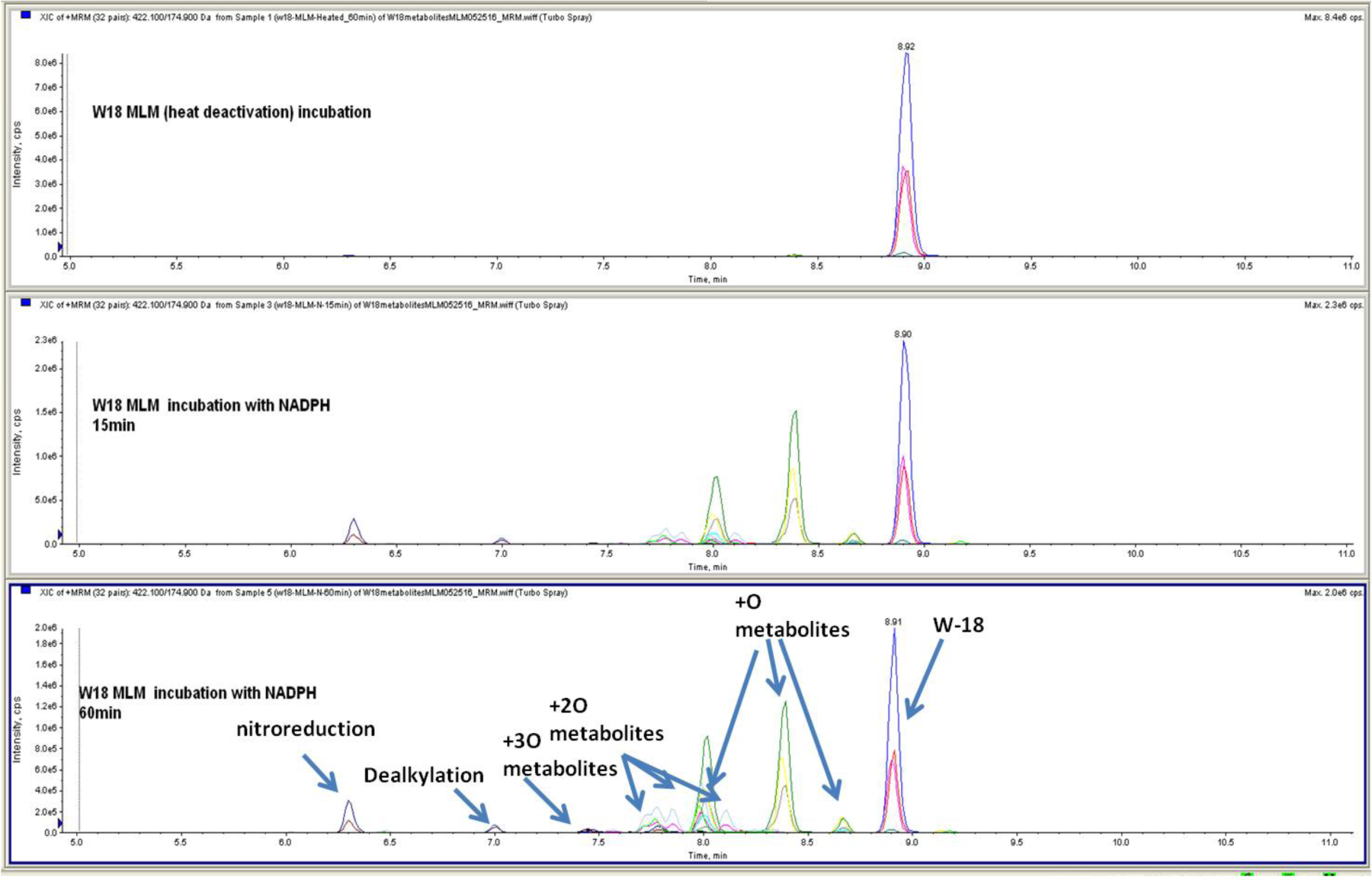
W-18 MLM incubations time points metabolities profiling analysis.

**Supplementary Figure 4.**
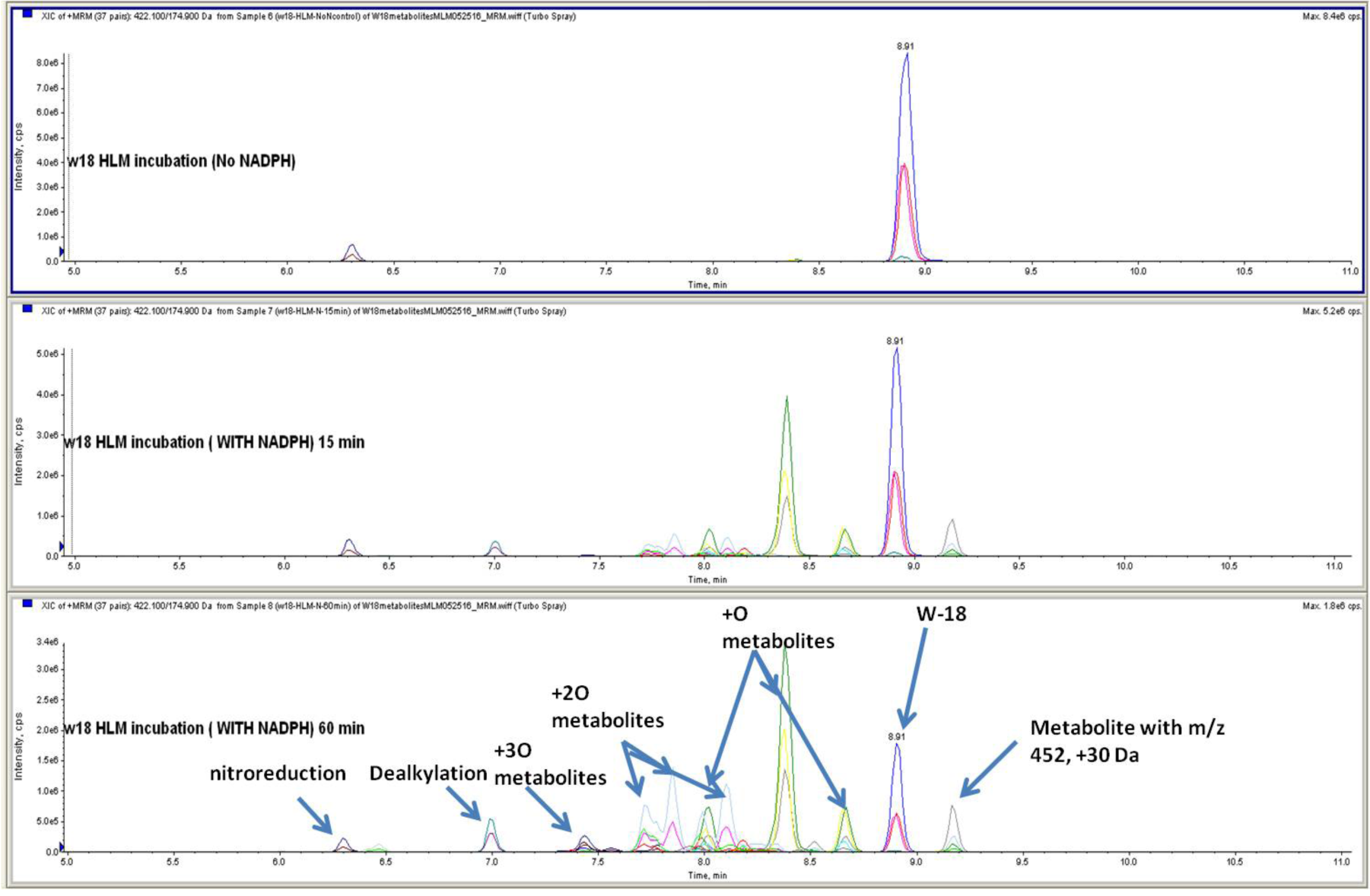
W-18 MLM incubations time points metabolities profiling analysis.

